# Spatially resolved mapping of histones reveal selective neuronal response in Rett syndrome

**DOI:** 10.1101/2025.06.23.661075

**Authors:** Frederike Schäfer, Guiseppa de Rocco, Moritz Völker-Albert, Ignasi Forne, Nicoletta Landsberger, Axel Imhof, Shibojyoti Lahiri

## Abstract

Rett Syndrome (RTT), a severe neurological disorder caused by loss-of-function mutations in the X-linked *MECP2* gene, results in profound life-long neurological dysfunction. RTT patients live an apparently normal initial life until 12-18 months of age following which, a progressive accumulation of a wide range of phenotypic manifestations sets in. While MeCP2 is known to regulate chromatin, its impact on global histone composition and dynamics remains poorly understood. Here, we combine mass spectrometry imaging (MSI) and laser capture microdissection (LCM) coupled to LC-MS/MS to systematically profile histone proteoforms in three key brain regions: the dentate gyrus (DG) and cornu ammonis (CA) of the hippocampus, and the cerebellum (Cb). Our analysis reveals striking neuron-specific differences in histone composition between Mecp2-deficient and wildtype (WT) mice. Interestingly, the expression of a pathogenic Mecp2 missense mutant (Y120D) results in subtler changes in histone composition that are distinct from the null mutations. This study provides the first spatially resolved epigenetic atlas of histone proteoforms in RTT and suggests that Mecp2 loss perturbs chromatin homeostasis in a neuron- and mutation-dependent manner. Our findings underscore the critical need for cell-type-resolved analyses to unravel the mechanistic underpinnings of RTT and emphasize the importance of personalised therapeutic strategies that consider both the affected cell-type and particular Mecp2 mutation.

## Introduction

Rett syndrome (RTT) is a disorder of the central nervous system (CNS) that primarily affects females and is the second most common cause for intellectual disability in females after Down’s syndrome^1^. Children with RTT typically exhibit apparently normal postnatal development during the first 12–18 months of life, after which a progressive neurological decline ensues. Hallmarked by escalating motor, speech, and cognitive impairments, the situation is further complicated by the progressive occurrence of several other symptoms such as scoliosis, seizures, respiratory dysrhythmias, ataxia, dysphagia and gastrointestinal problems, reflecting widespread and evolving neurological dysfunction ^2,3^.

RTT is caused by loss-of-function mutations in the X-linked gene encoding methyl-CpG-binding protein 2 (*MECP2*)^4,5^. Most of the missense mutations in *MECP2* that correlate with severity of symptoms have been found in either the methyl binding domain (MBD) and/or the transcription repression domain (TRD) ^2^. MeCP2 was initially identified as a transcriptional repressor that recruits co-repressor complexes containing histone de-acetylases (HDACs) ^6,7^. However, additional studies showed that MeCP2 can also affect gene expression through interactions with transcriptional activators ^8,9^. Beyond its role in the direct regulation of transcription, MeCP2 has also been reported to regulate global chromatin structure^8,10–14^. It induces chromocenter clustering^15^, maintains chromatin loops^16^ and nucleosome-free regions^17^ and compacts nucleosomal arrays both *in vitro* and *in vivo*^11,18,19^. Mecp2 is highly expressed in mature neurons where it partly replaces linker histone H1^20,21^. Loss of Mecp2 leads to an approximately 2-fold increase in linker histone H1 in neurons suggesting an overlapping set of chromatin associated functions^19,22^. It has been shown previously that loss of Mecp2 leads to diverse physiological effects depending on the type of affected neurons^23–30^. There are two different yet non-exclusive hypotheses for why a specific knockout results in different physiological outputs. On one hand, deletion of the *Mecp2* gene may disrupt a core function shared across neurons, with phenotypic differences arising simply from the distinct roles of the affected brain regions. On the other hand, cell-type-specific studies reveal that Mecp2-deficiency alters divers gene sets in different neuronal populations suggesting that Mecp2’s molecular role is highly context-dependent. Resolving these models requires spatial analysis of molecular changes upon *Mecp2* removal *in situ* to investigate whether regional phenotypes stem from uniform dysfunction or cell-type-specific mechanisms. In addition, expression of *Mecp2* is spatio-temporally regulated making it crucial to study the effects of its loss in the spatial context^31,32^.

Histone proteoforms—including canonical histones, their variants, and post-translationally modified forms—play a critical role in regulating chromatin architecture and shaping the epigenetic landscape in neurons^33–39^. As a result, they have been associated with or implicated in several neurodegenerative diseases^40–44^. Mecp2, as a chromatin-associated protein, is widely believed to influence these structural and regulatory processes^10–12,17,39,45^. Therefore, we wanted to comprehensively study the effect of Mecp2 loss-of-function on the histone composition *in situ*. So far, there have been few studies investigating the effects of Mecp2-loss on histone proteoforms. These studies, however, were restricted by the availability of specific antibodies, lack of spatial context and a comprehensive understanding of the entire histone proteoforms landscape^45–49^. To address this gap we used a dual spatial approach that involves mass spectrometry imaging (MSI) and laser capture microdissection (LCM) coupled to LC-MS/MS. This strategy enabled unbiased, high-resolution quantification of histone proteoforms across distinct brain regions. Based on preliminary observations and physiological context, we focussed our measurements on the dentate gyrus granule cell layer and cornu ammonis pyramidal neurons (CA) of the hippocampus, and the cerebellar granule cell layer (Cb).

Mecp2-deficiency induces profound, cell type-specific alterations in nucleosome composition, total histone levels and modification patterns. We observe that the levels of all histone variants including those of linker histone H1 are substantially disturbed in absence of Mecp2. In addition, lack of Mecp2 is also associated with major changes of global modification patterns across the neuronal subpopulations. These changes occur in the absence of significant alterations in the expression levels of histone-modifying enzymes, implicating post-translational mechanisms or enzyme redistribution as key drivers of the observed chromatin remodelling. We compared the findings with neuronal populations of a mouse model carrying the human pathogenic mutation Y120D^50^. The *Mecp2^Y120D/y^* model (KI) recapitulates RTT pathology and exhibits distinct chromatin dynamics to those observed in Mecp2 knockouts (KO) ^50,51^. We observe that the effects on histone proteoforms in the knock-in model is discordant to those of the knock-out, indicating at the possibility of distinct molecular pathways underlying different mutations for RTT.

Taken together, our work provides a spatially resolved atlas of histone proteoforms in wildtype mice and their alterations in RTT. We highlight the importance of analysing chromatin in a spatially resolved manner. By linking Mecp2 loss-of-function (both in null and mutant models) to discrete histone proteoform alterations, we provide novel insights into the possible epigenetic mechanisms driving RTT associated with different Mecp2 mutations.

## 2 Methods

### 2.1 Animals

Both mouse lines were housed in the animal facility of the San Raffaele Scientific Institute of Milan. The *Mecp2* null strain was originally purchased from Jackson Laboratories (B6.129P2(C)-Mecp2tm1.1Bird/J) and transferred on a CD1 genetic background^52^. The *Mecp2^Y120D^* mouse strain was generated as reported in Gandaglia et al., 2019^50^. Both lines are maintained on a clean CD1 background by crossing *Mecp2^-/+^*or *Mecp2Y^120D/+^* heterozygous females with WT CD1 males purchased from Charles River Laboratories. Mouse genotype was determined by PCR as previously described^50,52^. Mice were housed in groups of five in Tecniplast cages, on a 12h light/dark cycle in a temperature-controlled environment (21 ± 2°C) with food and water provided ad libitum. All procedures were performed in accordance with the European Community Council Directive 2010/63/UE for care and use of experimental animals; all the protocols were approved by the Italian Minister for Scientific Research and by the San Raffaele Scientific Institutional Animal Care and Use Committee in accordance with the Italian law.

Whole fresh brain were obtained from P42/45 wildtype or mutant littermates sacrificed by rapid decapitation.

### 2.2 MALDI-Imaging Mass Spectrometry

#### 2.2.1 Sample preparation

Three mouse brains of each genotype were halved mid-sagitally and sectioned on a Leica CM3000 cryostat (Leica Biosystems, Germany). 12*µ*m sections were thaw-mounted onto indium-tin-oxide (ITO) slides (Bruker Daltonics GmbH, Germany) coated with polyL-Lysine (1:1 in water, 0.1% NP40). Sections were stored in a sealed container at -80^◦^C until usage.

Sections were washed according to a preestablished protocol to remove interfering salts and lipids^53^. In brief, proteins were precipitated *in situ* by incubating in 70% Ethanol and 100% Ethanol for 30s each, followed by 2min incubation in Carnoy’s fluid (60% EtOH, 30% chloroform, 10% acetic acid (v/v/v)). The slides were dried under vacuum overnight at room temperature. Sections were then sprayed with matrix (10 mg/ml sinapinic acid (SA) in 60% ACN; 0.2% TFA) using a HTX TM Sprayer (HTX Imaging, HTX Technologies) with the standard manufacturer’s method.

#### 2.2.2 MALDI-Imaging mass spectrometry

Tissue measurements were carried out using a rapifleX MALDI tissuetyper MALDI-TOF/TOF mass spectrometer (Bruker Daltonics GmbH, Germany). Samples were measured with 50*µ*m spatial resolution and within a mass range of 500-3.200 m/z using positive reflector mode. The instrument was calibrated prior to measurement using a peptide calibrant solution (Protein calibration standard I, Bruker Daltonics GmbH, Germany) spotted onto the same slide.

#### 2.2.3 Histology and Nuclear density determination

After MSI experiments, the matrix was removed and the tissue was stained with hematoxylin and eosin (HE). Nuclear density was determined as previously described^54^. In brief, slides were scanned with a Mirax Desk digital slide scanner (Carl Zeiss MicroImaging, Germany) at 20x objective magnification. Images of HE stained slides were analyzed using Definiens TissueStudio3 (Definiens AD, Germany). Nuclei were identified and quantified based on morphology, size, pattern and neighborhood. The generated HE images were also used for co-registration to IMS data to allow histological correlation.

#### 2.2.4 Data analysis

IMS data was TIC normalized using fleXImaging (Bruker Daltonics GmbH, Germany). Regions of interest were manually selected based on correlation with morphological staining. Neuronal populations of dentate gyrus granule cells, cornu ammonis pyramidal cells and cerebellar granule cells were chosen. Regional mass lists were exported, peak picking was carried out and masses were matched to theoretical histone masses. The IMS data files were imported into SCiLS Lab 2016b (Bruker Daltonics, www.scilslab.de) and unbiased hierarchical clustering was performed using the segmentation pipeline with a minimal interval of 5 Da^55^.

### 2.3 LC-MS/MS

#### 2.3.1 Laser Capture Microdissection

Three mouse brains were cryosectioned for *Mecp2^-/y^* and their respective CD1 WT and *Mecp2^Y120D/y^*. Due to limited availability, two replicates were used for Mecp2^Y120D/y^ WT littermates. 50 *µ*m sagittal sections were produced and mounted onto metal-frame PET slides (Leica Biosystems, Germany). The sections were stained with Hematoxylin and Eosin (HE) staining to visualize brain regions for LCM. A Leica LMD 7000 with a 5x objective was used for LCM of brain regions. The laser setup was configured as follows: Pulse frequency - 120, maximum pulse energy - 50, aperture - 17, Speed - 10, Head current - 100 %, Offset - 65. Hippocampal neuronal layers dentate gyrus granule cells and cornu ammonis pyramidal cells (CA) as well as the cerebellar granule cell layer (Cb) were dissected. Excisions of ten serial sections were pooled for each biological replicate and sectioned areas traced for post-measurement normalization. Separate sample pools of ten sections each were prepared for total proteome, histone and DNA extraction.

#### 2.3.2 Acidic extraction of histones

Laser-microdissected samples were processed using an in-house protocol for histone extraction based on previous work^56^. In brief, lysis buffer (60mM KCl, 15mM NaCL, 4mM MgCl_2_, 15 mM HEPES pH=7.6, 0.5% Triton-X, 1mM DTT, 1 tablet Roche cOmplete protease inhibitor) was added to the samples and sections were manually homogenized using Micro-homogenizers, PP (Carl Roth GmbH+Co. KG) to enhance tissue lysis. The samples were centrifuged and the resulting pellet containing the nuclei resuspended in 0.2M H_2_SO_4_ and incubated overnight at 4^°^C. Basic proteins were precipitated with trichloroacetic acid (TCA). Lysine residues were acylated with NHS-propionate prior to tryptic digestion. Stable-isotope labeled (SIL) histone peptides (label: R(15N; 13C); 500fmol/*µ*l; JPT Peptide Technologies GmbH, Germany) were added to the digestion mix to serve as internal control for identification^57^. Samples were digested at 37^°^C overnight with trypsin. Tryptic peptides were desalted and purified using C18 Stagetips. The samples were then dried in a vacuum desiccator and resuspended in 10µl loading buffer (0.3% TFA, 2% ACN) for LC-MS/MS measurement.

#### 2.3.3 DNA purification and measurement

Genomic DNA was purified from laser-microdissected tissue using the NucleoSpin Tissue Kit (MachereyNagel GmbH&Co. KG). To enhance tissue lysis, tissues were initially homogenized with Microhomogenizers, PP (Carl Roth GmbH+Co. KG). DNA concentration was determined using the Qubit dsDNA High Sensitivity Assay Kit (Thermo Fisher Scientific Inc.).

#### 2.3.4 Proteomics sample preparation

Laser-capture microdissected tissue was processed using the PreOmics iST Kit for mammalian tissue according to the manufacturers guide (PreOmics GmbH). Tissue lysis was enhanced by manual homogenization with Micro-homogenizers PP (Carl Roth GmbH+Co. KG) after the addition of lysis buffer. Protein digestion with trypsin/LysC was carried out for 3h at 37^°^C. Resulting peptides were desalted, purified and dried in a speedvac, followed by resuspension in 12 µl “LC-LOAD” for LC-MS/MS measurement.

#### 2.3.5 LC-MS/MS

For LC-MS/MS, the peptides were injected into an Ultimate 3000 RSLCnano system and separated either in a 25-cm analytical column (75 µm ID, 1.6 µm C18, Aurora-IonOpticks) with a 50-minute gradient from 2 to 35% acetonitrile in 0.1% formic acid. The effluent from the HPLC was electrosprayed directly either into an Orbitrap QExactive-HF (Proteome analysis) or an Orbitrap Exploris 480 (Histone extraction) instrument both operating in data-dependent mode to automatically transition between full-scan mass spectrometry and MS/MS acquisition. Typical mass spectrometric conditions for both instruments were: spray voltage, 1.5 kV; no sheath and auxiliary gas flow; heated capillary temperature, 275°C; ion selection threshold, 5×10^3^ counts; dynamic exclusion, 20s). Specific settings are described below. QExactive-HF settings: Survey full-scan MS spectra (from m/z 375–1600) were acquired with resolution R=60,000 at m/z 400 (max IT 60 ms, AGC target of 3×10^6^). The 10 most intense peptide ions with charge states between 2 and 5 were sequentially isolated to a target value of 1×10^5^, fragmented at 27% normalised collision energy and acquired with resolution 15000 at m/z 400 (max IT 60 ms). Exploris 480 settings: Survey full scan MS spectra (m/z 250–1200) were acquired with resolution 60000 at m/z 400 (max IT 40 ms, AGC target of 3×10^6^). The 15 most intense peptide ions with charge states between 2 and 6 were sequentially isolated to a target value of 2×10^5^, fragmented at 30% normalized collision energy and acquired with resolution 15000 at m/z 400 (max IT 40 ms).

#### 2.3.6 Data analysis

MaxQuant 2.0.1.0. was employed for protein identification. Default MaxQuant conditions were used with parent and fragment ion mass tolerances of 20 ppm. Allowance of missed cleavages was set to 2. For the searches the reviewed mouse canonical protein database from Uniprot was used. Additionally, the following conditions were used: protein FDR and peptide FDR of 0.05 and 0.01, respectively; minimum peptide length of 5; variable modifications of oxidation (M), acetyl (protein N-term), acetyl (K), methyl(KR), dimethyl (KR) and trimethyl (KR); variable modification of carbamidomethyl (C); peptides for protein quantification, razor and unique; minimum peptides of 2; minimum ratio count of 2. Protein identifiation was carried out based on a unique peptide detected in all or at least 50% of the replicates.

Due to the high sequence similarity between histone family members, quantification was carried out on the peptide-level to ensure accurate evaluation. One shared peptide was used to approximate total histone abundance. Peptides were chosen based on the following criteria: maximum completeness of detection across samples, identification by MS/MS and maximum coverage ensured by peptide length. Likewise, histone isoforms and variants were quantified based on a unique peptide that was chosen based on the same criteria. As H1.0 does not share any peptides with H1.1-1.5, a separate peptide was used for H1.0 and intensities summed up to estimate the total H1 pool. Peptide sequences can be found in Supplemental File S2. Peptide intensities were normalized to the DNA concentration of the respective sample, batch corrected using ComBat^58^ and imputed using Perseus 2.0.6.0 ^59^ where necessary.

Post-translational modifications of H3 and H4 peptides were manually validated using Skyline^60^. Identification was guided by spike-in internal standards. Modified peptides had to be present across samples to be considered for analysis. Intensities were normalized to DNA concentration and batch corrected using ComBat^58^. The relative abundance of each modified peptide was calculated as percentage of the overall peptide according to previous work^56^. For fold change comparison of absolute value of modified peptides between WT and mutant, modified H3 peptides were normalized to unmodified H3_41-49 peptide and modified H4 peptides were normalized to unmodified H4_46-57 peptide.

Statistical significance was probed using a two-tailed t-test. Results can be found in Supplemental File S5.

## Results

### Spatially resolved quantification of histone proteoforms

To investigate Mecp2 loss-of-function–induced molecular alterations in the brain *in situ*, we employed Mass Spectrometry Imaging (MSI) to spatially resolve intact proteoforms in an unbiased, high-throughput manner (Fig. 1A). This approach enabled a proteome-wide assessment of regional molecular signatures without prior target selection. Unsupervised clustering of MSI signals revealed that hippocampal neurons (DG and CA regions), cortex, and cerebellar granule layer (Cb) formed distinct spatial clusters early in the analysis (Fig. 1B; see dendrogram). Principal Component Analysis (PCA) confirmed the clear separation among the exclusive neuronal populations (Fig. 1C), primarily driven by strong histone signals (Fig. S1A; Supplemental file S1, sheet: Loadings plot) as a result of high nuclear density^54^. While the overall clustering patterns appeared similar between Mecp2-deficient and WT brains, we observed pronounced differences in the composition of histone proteoforms (Fig. 1D, Supplemental file S1, Sheet: MSI histone masses). Notably, the clear separation of cerebellar and hippocampal regions seen in WT brains was disrupted in their RTT counterparts, indicating altered composition of histones in these areas. Given the known physiological relevance of these brain regions to RTT pathogenesis (Fig. S1B), we micro-dissected them using laser capture (LCM) and performed LC-MS/MS analysis for high-resolution, exhaustive and precise identification and quantification of histone proteoforms (Fig. 1E). This integrative spatial-proteomic strategy allowed us to resolve the region-specific histone landscape potentially underlying the RTT phenotype. Histones were acid extracted and subsequently analysed using a tailored protocol for quantitation of histone variants and modified residues (Fig. 1E). Based on the extracted ‘proteome’, our regions of interest clustered separately within the principal component space, with the hippocampal populations closer together than the cerebellum (Fig. 1F). Our protocol was sensitive enough to quantify more than 30 different variant-specific as well as modified peptides from a minimum of 30,000-35,000 cells (from CA). We identified a minimum of 11 different core and linker histone isoforms or variants and profiled 20 modifications on H3/H4 in WT and RTT brains (Fig. 1G).

**Figure 1:**
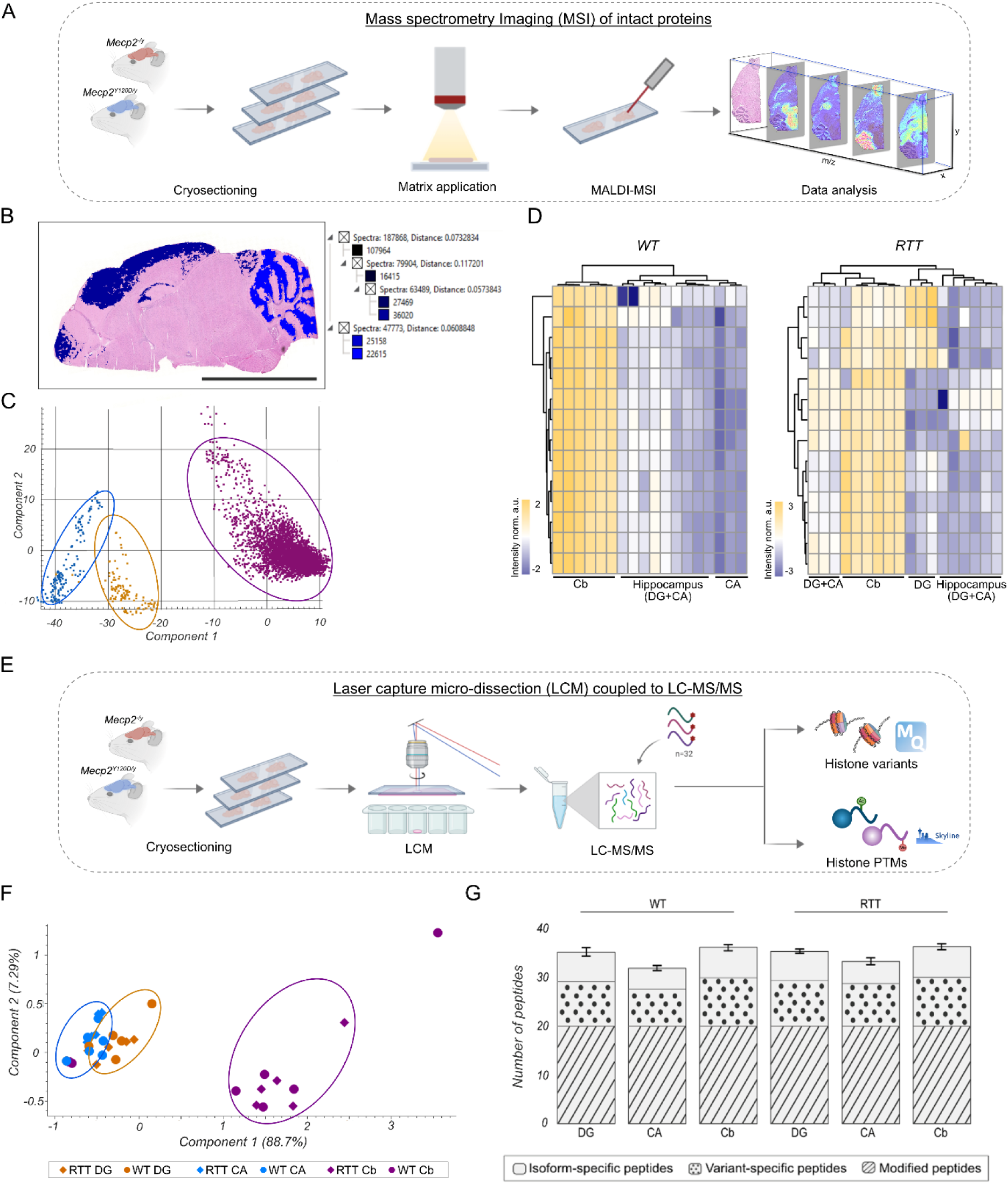
Rapid spatial screening reveals altered histone proteoforms. **A.** Workflow for MALDI-MSI. After cryosectioning, brain sections are sprayed with matrix and analysed using rapifleX tissuetyper, followed by data analysis in flexImaging and SCiLS Lab. **B.** Hierarchical clustering of MSI data reveals early separation of cortex/hippocampus (darkblue) and cerebellum (blue). Dendrogram of unbiased clustering analysis. Scale bar = 5mm. **C.** PCA of MSI data. Dots represent individual pixels. Neuronal populations of CA (blue), DG (orange) and Cb (pink) can be separated based on their proteomic signature generated by MALDI-MSI. **D.** Unbiased clustering of MSI-detected histone proteoforms in WT and RTT model brains. Histone identities were assigned to masses based on previous work^50^. Each column represents a biological replicate of WT, *Mecp2^-/y^*and *Mecp2^Y120D/y^*. **E.** Workflow for laser capture microdissection (LCM) coupled to LC-MS/MS. Mouse brains were cryosectioned and regions of interest excised by LCM. Histone extraction was performed on collected tissue samples with the addition of 32 stable isotope labelled (SIL) peptides. Intact histones were identified and quantified by MaxQuant and modified peptides were manually picked and quantified in Skyline. **F.** PCA of LC-MS/MS data of basic proteins. These proteomic signatures are sufficient to separate the neuronal population DG, CA and Cb. **G.** Number of detected peptides of canonical histone isoforms, variants and modified peptides are reproducible across sample groups with > 30 proteoforms.

### Loss of Mecp2 induces variance in histone levels

#### Canonical histones

Our approach allowed us to precisely quantify the amount of histones in WT and *Mecp2^−/y^* brains. To quantify the total level of core histones we used peptides that were detected in all replicates and all respective isoforms (40-49 for H3, 47-56 for H4, 22-30 for H2A and 101-121 for H2B). To account for potential differences in genomic material that could possibly influence histone amounts, intensity of the individual peptides where normalized to the total amount of DNA present in the respective samples (Fig. S1C). Looking at region-specific histone alterations, we observe opposing trends within the neuronal populations of the hippocampus (Fig. 2A). While DG shows a moderate decrease in of all core histones levels, the CA is associated with minor upregulation of the same. Strikingly, cerebellar granule cells show an imbalance of histone amounts with an increase of H3 and H2B and a reduction in H2A and H4 (Fig. 2A). This disparity in histone stoichiometry might possibly indicate towards a fraction of unbound histones within the cell.

**Figure 2:**
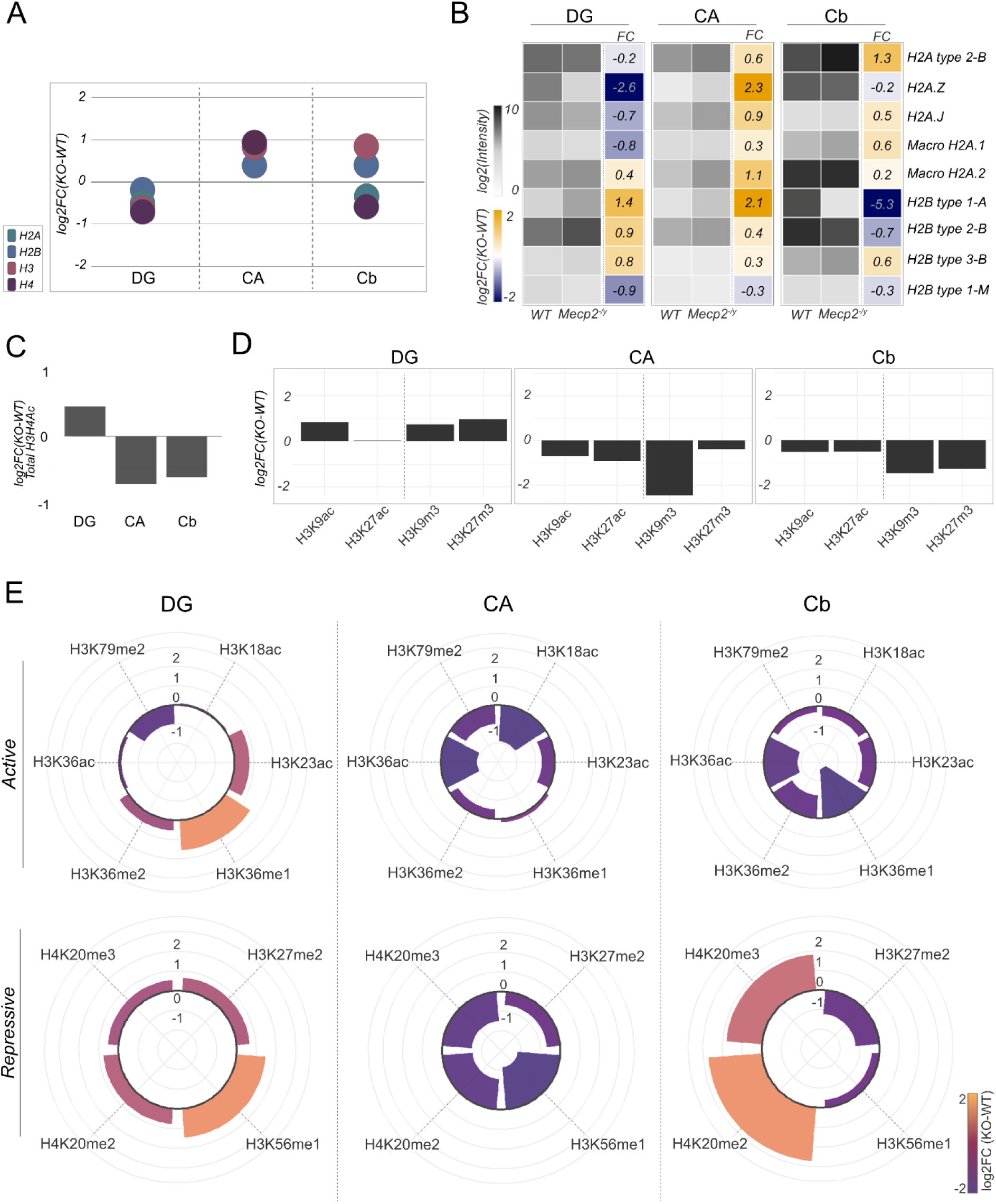
Neuronal histone proteoform landscape is altered by loss of *Mecp2*. **A.** Alteration of total histone abundance upon loss of *Mecp2* in neuronal populations DG, CA and Cb in log2 fold change. n=2-3 **B.** log2 intensities of H2A/B peptides in WT and *Mecp2^-/^*^y^ brains and respective **f**old changes in DG, CA and Cb. **C.** Fold change of total acetylation of H3/H4. **D.** Fold change of H3K9/K27 acetylation (ac) and tri-methylation (me3) in neuronal populations DG, CA and Cb. **E.** Fold change of histone post-translational modifications associated with active and repressive transcription.

#### Histone variants and isoforms

The core histone proteins are also characterised by the presence of variants that have distinct functional roles including regulation of neuronal functionality^33,34,36–38^. Therefore, change of relative proportions of these variants might play a crucial role in brain functionalities, particularly in the absence of key chromatin modulator Mecp2. Histone H3.3 is one of the most important histone variants in the adult mammalian brain, as it constitutes majority of the H3 pool^33,34,61^. Similar to previous findings, we observe that H3.3 makes up the majority (50-60%) of the total H3 pool in hippocampal neuronal populations, while the contribution is lower in Cb^34^. This relative proportion of the H3.3 variant remains unaffected in granule layers of Mecp2 null mice but is reduced in pyramidal neurons (Fig. S2C; Fig. S3A)

All H2A/B variants and isoforms were quantified based on corresponding unique peptides (Fig. S2B; Supplemental File S2) detected across most samples. Expectedly, WT brains are characterized by high expression of the canonical H2A type 2-B and macroH2A.2 across all neuronal populations^62^ (Fig. 2B). In addition, granule layers of DG and Cb show high abundance of H2A.Z while relative contribution of H2A.J to the total H2A pool is more dominant within the hippocampus as compared to the Cb (Fig. 2B). Upon loss of Mecp2, different variants and isoforms are altered in a cell-type specific manner (Fig. 2B). Most noteworthy is the strong up - and downregulation of H2A.Z in CA and DG, respectively, followed by H2A.J and macroH2A.1, albeit to a lesser extent. Except for macroH2A.2, all other H2A variants show an opposite trend of dysregulation within the two hippocampal neuron populations (Fig. 2B). In the *Mecp2^-/y^* Cb, however, H2A.Z stands out in being slightly downregulated as compared to all other H2A variants (Fig. 2B). While canonical H2A remains the most abundant isoform in KO neurons, relative contribution of variants is distinct to that of WT, suggesting neuron-specific compositional changes of the cellular H2A pool (Fig. 2B).

For H2B, we primarily detect canonical H2B isoforms that are located either within the major histone cluster on chromosome 13 (H2B type 1A and H2B type 1M) or on the minor clusters on chromosome 3 (H2B type 3B) or 11 (H2B type 2B) (Fig. 2B)^63,64^. Similar to H2A, loss of Mecp2 induces regional reorganisation of the H2B pool, as indicated by distinct alterations and relative abundances of H2B isoforms across different regions of the *Mecp2^-/y^* brains (Fig. 2B).

#### Core histone modifications

Mecp2 is known to co-recruit histone deacetylases (HDACs) as part of a repressor complex, yet reports on its impact on global histone acetylation are inconsistent^22,47^. While some studies show increased H3 acetylation in neuronal nuclei from mouse brain^22^, others report no change in clonal cell cultures from RTT patients, which instead exhibit hyperacetylation of H4^47^. To resolve these apparent discrepancies, we aimed to systematically analyse total histone acetylation in a cell-type–specific manner within the RTT brains. In *Mecp2^-/y^* brains, we observed only DG granule cells to be associated with an overall increase in total H3/H4 acetylation, whereas the other two investigated cell populations show a global depletion of the acetylation mark (Fig. 2C). This does not seem to be facilitated by altered levels of Mecp2-recruited modifiers, as we do not see changes in overall HDAC abundance (Fig. S2F).

Two most prominent class of active and repressive marks studied in the context of Mecp2-chromatin interaction are the tri-methylation (me3) and acetylation (ac) of the K9 and K27 residues of core histone H3^45,48,49^. However, the impact of Mecp2 loss on these epigenetic modifications remains unclear, particularly globally and in terms of cell-type–specific patterns that may underlie the pathophysiology of Rett syndrome.

Our unbiased approach enabled us to simultaneously profile these modifications in the neuronal populations of interest (Supplemental File S3). WT and *Mecp2^-/y^* brains display a unique histone modification landscape, independent of total histone levels. The relative proportion of individual marks to the overall peptide abundance differ substantially (Fig. S2D). This gives valuable insight into contributions of modified peptides within a genotype, yet it does not portray absolute levels. As histone amounts themselves are subjected to change, modification levels have to be corrected for these altered abundances and directly compared between genotypes. Here, we observe that the trends of dysregulation of these modifications follow a cell-type specific pattern (Fig. 2D). Upon loss of Mecp2, both the acetylation (ac) and trimethylation (me3) marks of H3K9 and H3K27 are enriched in the DG, whereas they are depleted in the other two neuronal populations, albeit to quite different extents (Fig. 2D). These findings likely reflect the ambivalent nature of Mecp2 function, having both repressive and activating effects on local gene expression. Our approach captures an accumulation of local dysregulation of histone proteoforms across different neuronal populations, revealing cell-type–specific selectivity as an additional layer of epigenetic dysregulation in RTT beyond gene-specific alterations.

Additionally, we could profile many other active and repressive histone marks (outside of H3K9/27 me3/ac). In *Mecp2^−/y^* DG, the chromatin is characterized by an enrichment of active gene expression marks such as H3K23ac and H3K36me1/2 along with a global gain of repressive marks (Fig. 2E). However, the CA and Cb exhibit an overall depletion of these marks except for H3K36me1 in CA and H4K20me2/3 in Cb (Fig. 2E). Notably, these changes of modified peptides are anti-correlated to changes in total H3/H4 levels, indicating preferential gain/loss of unmodified proteoforms (Fig. 2A; Fig. S2E).

The observed alterations in the chromatin landscape, as reflected by changes in histone modifications, are not facilitated by differential regulation of the corresponding modifiers, which either remain consistent across cell types or the direction of dysregulation doesn’t align with the effect of the modification it drives (Fig. S2F). This suggests that the dysregulation arises most likely from altered enzymatic activity and/or impaired recruitment to chromatin.

Taken together, loss of Mecp2 function induces complex changes in the histone modification landscape that are unique for specific neuronal populations. The observed alterations likely reflect local changes in chromatin state that vary in magnitude and pattern within each neuronal subtype, highlighting the context-dependent nature of epigenetic dysregulation in *Mecp2^-/y^*brains.

#### Alterations of linker H1 levels and variant composition

Elevated histone H1 expression is regarded as a potential compensation for Mecp2 loss-of-function due to its increased deposition on chromatin in neurons of *Mecp2^−/y^* mice. However, the underlying mechanisms and its generality remain elusive due to conflicting evidence^19–22^.

Our approach allowed us to profile the total H1, including all H1 variants, comprehensively across the neuronal populations. For accurate quantification, one shared peptide among all H1 variants was taken into consideration to approximate for total H1 levels. We indeed observe an increase of total H1 in Cb, whereas the two hippocampal neuronal populations show different trends (Fig. 3A). While levels show a slight increasing trend in CA, total H1 levels are reduced in *Mecp2* KO DG (Fig.3A). Considering that mammalian H1 has 7 somatic variants and relative expression of these variants possibly vary between different cell types^65,66^, we attempted to understand dynamics of H1 incorporation upon Mecp2 loss-of-function. The applied methodologies allow us to quantify the relative contribution of different H1 variants to the total H1 pool based on one unique peptide characteristic to each variant that was detected across samples. Information about the unique peptides and their respective positions within the protein are provided in Fig. S2A and Supplemental File S2. As expected, H1.4 and H1.0 are the predominant H1 variants across all WT neuronal populations^65^, with the addition of H1.3 in granule cells (Fig. 3B). These variants are affected differently by loss of Mecp2, resulting in an altered composition of the H1 pool in *Mecp2^-/y^* brains. Overall, within the hippocampal populations of *Mecp2* KO, we observe that most H1 variants show a decrease in DG, while the trends are opposite in CA with the exception of H1.5 (Fig. 3B). Nonetheless, H1.0, H1.3 and H1.4 remain the variants with the most prominent contribution to the total H1 pool, despite different absolute levels compared to WT (Fig. 3B). A mixed effect characterises the Cb, where the replication-dependent variants H1.1-H1.5 increase while H1.0 is depleted (Fig. 3B). This further reduces the proportion of H1.0 to the H1 pool in Cb, while H1.1 and H1.2 now substantially contribute to the pool as compared to the WT (Fig. 3B). This reflects the cell-type specific contribution of individual somatic H1 variants and suggests against inferring just from total H1 levels.

**Figure 3:**
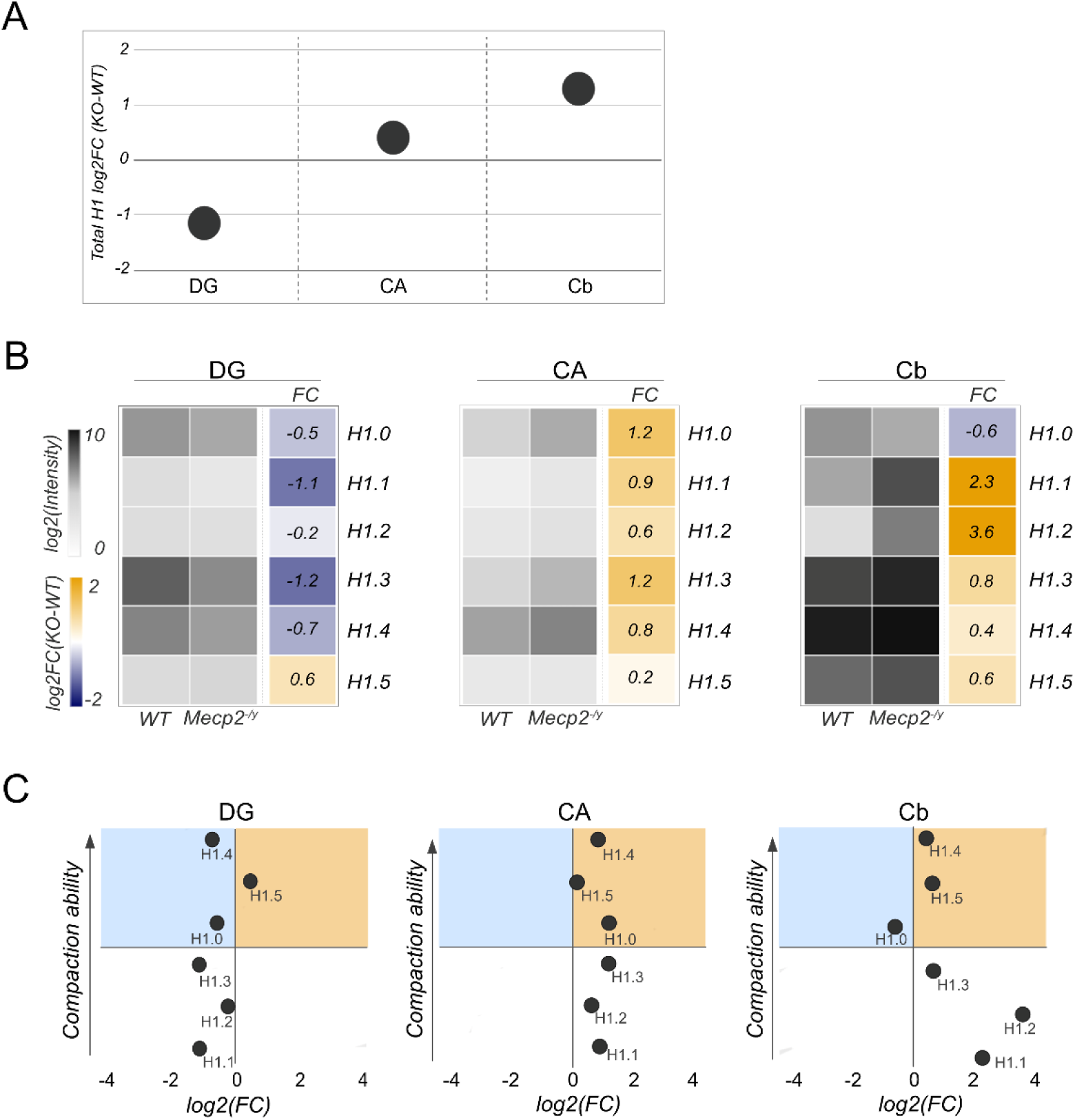
Histone H1 is affected by *Mecp2* loss in a neuron-specific manner. **A.** Total abundance of H1 in WT and *Mecp2^-/y^* in log2 fold change. n=2-3. **B.** log2 intensities of H1 variant peptides in WT and *Mecp2^-/y^* brains and respective **f**old changes in DG, CA and Cb. **C.** Fold changes of histone H1 in correlation to their compaction ability. Compaction ability was scored based on previous classification by Prendergast et al. Positive (orange) and negative (blue) fold changes of variants with high compaction ability are highlighted.

The degree of influence of different H1 variants on chromatin compaction depends on their differential chromatin binding affinity and compaction ability^65^. Although the relative variant contribution to the total H1 pool might be different, the fold change of individual variants between the WT and *Mecp2^-/y^* brains provides an indication about possible H1-mediated chromatin compaction states (Supplemental File S4). In *Mecp2^−/y^* DG, variants with a higher compaction ability are not particularly enriched (Fig. 3C, left panel). The strongest compactor H1.4 shows an enrichment in CA along with substantial enrichment of variants having a moderate compaction ability (Fig. 3C, middle panel). However, in Cb, the enrichment of H1 variants include both the strongest compactors in the form of H1.4 and variants (H1.1 and H1.2) with lower compaction ability (Fig. 3C, right panel).

### 3.4 Effects on histone proteoforms are mutant-specific

Comprehensive characterisation of histone proteoforms in the *Mecp2* KO mice reveals that loss-of-function of Mecp2 has a profound and cell-type specific effect on the histone composition. However, total absence of Mecp2 might lead to compensatory mechanisms within the brain. We therefore characterised a mouse model harbouring the pathogenic mutant *Mecp2^Y120D/y^*that has previously been shown to have discordant chromatin features in spite of having similar phenotype ^50^. Unlike *Mecp2^-/y^* animals, total histone levels change only marginally in the hippocampus (DG and CA), while cerebellar granule cells display an increase of H2A/B (Fig. 4A). With respect to core histone variants, we observe that the effects have a cell-type specific signature that is different from the KO animals (Fig. 4B, Fig. 2B). Wildtype mice carry the canonical H2A type 2-B as the major H2A isoform, followed by macroH2A.2 and H2A.J (Fig. 4B). Unlike the KO animals, contribution of H2A.Z is considerably enriched in *Mecp2^Y120D/y^* hippocampus, whereas it is unchanged in the Cb (Fig. 4B). Interestingly, as opposed to *Mecp2^-/y^* Cb, the knock-in animals display a marked depletion of both macroH2A.2 and H2A.J. This is accompanied by a substantial enrichment of the canonical H2A type 2-B contribution, resulting in a notable reduction in variety of the H2A pool (Fig. 4B). H2B isoform expression and contribution differs between WT and *Mecp2* KI mice, displaying isoform-specific alterations across all neuronal populations (Fig. 4B) that are different from its KO counterparts (Fig. 2B).

**Figure 4:**
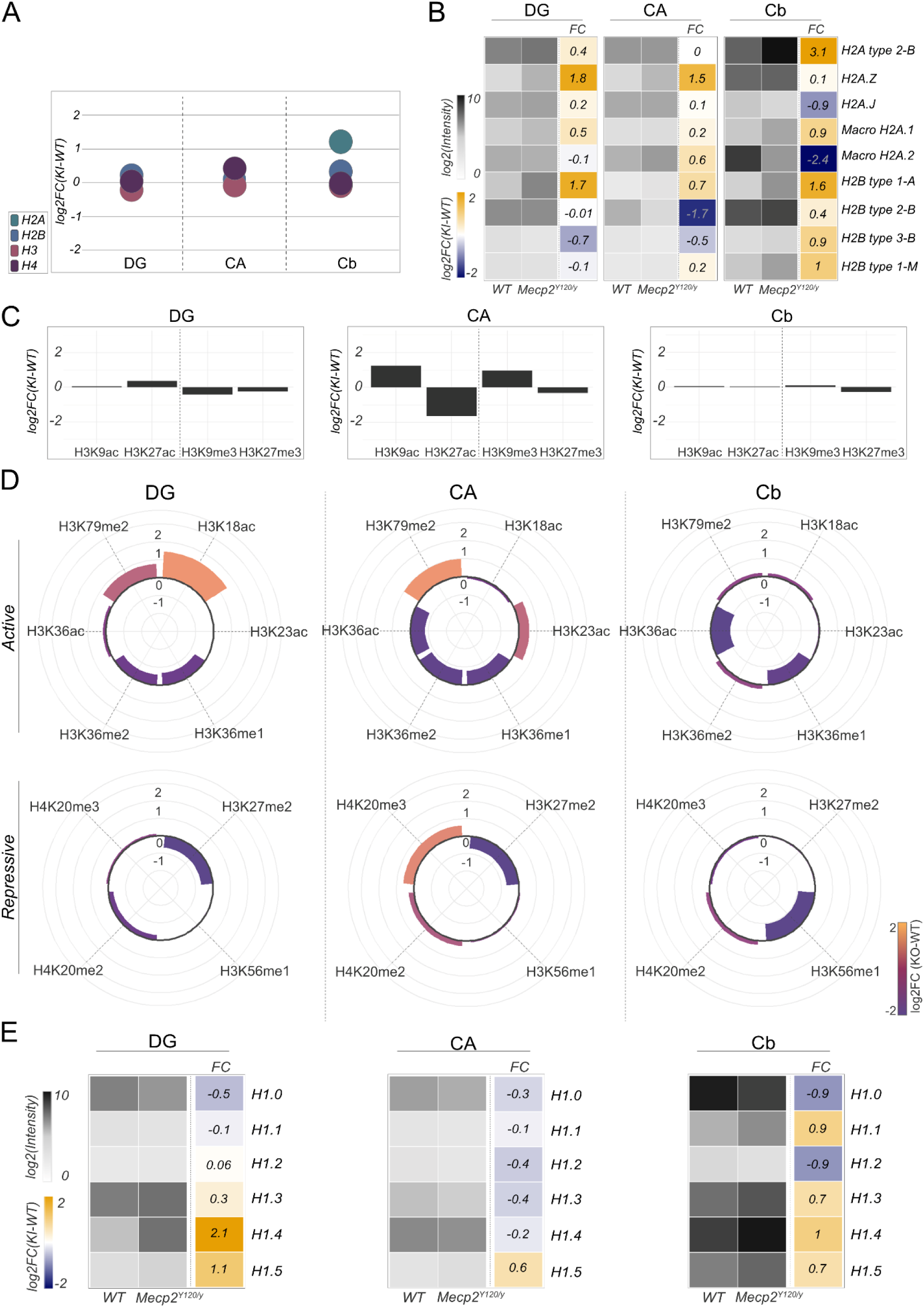
*Mecp2^Y120D/y^* induces distinct changes in neuronal histone proteoform levels. **A.** Alteration of total histone abundance in neuronal populations DG, CA and Cb of *MeCP2^Y120D/y^* brains. **B.** log2 intensities of H2A/B peptides in WT and *Mecp2^Y120D/y^* brains and respective **f**old changes in DG, CA and Cb. **C.** Fold change of H3K9/K27 acetylation (ac) and tri-methylation (me3) in neuronal populations DG, CA and Cb. D. Fold changes of histone post-translational modifications associated with active and repressive transcription show unique trends across the cell types. **E.** log2 intensities of H1 variant peptides in WT and *Mecp2^Y120D/y^* brains and respective **f**old changes in DG, CA and Cb.

The Y120D mutation also has distinct effects on the deposition of histone modifications. Global H3/H4 acetylation levels are largely unchanged in DG, while they show subtle increase in CA and a decrease in Cb and therefore differ substantially from KO brains (Fig. S3B).

The histone modification landscape of the WT littermates is comparable to *Mecp2^-/y^* WT littermates, despite minor deviations in relative abundance of single modified peptides. However, in mutant animals the composition of modified peptides is notably different (Fig. S3D, S2D). *Mecp2^Y120D/y^* CA and Cb only show minor changes in relative abundance of modified peptides. DG on the other hand, shows a particularly high proportion of acetylated histone H3 peptides (Fig. S3D).

These minute compositional changes reflect slight changes in absolute levels. Here, the presence of the mutant protein has minimal effects on H3K9 and H3K27 tri-methylation (me3) and acetylation (ac) in granule cell layers of both DG and Cb (Fig. 4C). In CA, the marks on H3K9 are enriched while H3K27 are depleted, particularly affecting H3K27ac (Fig. 4C) and deviating substantially from the situation in *Mecp2^-/y^* brains. Global effects on other active and repressive histone modifications show a cell-type specific signature that is distinct from *Mecp2^-/y^* brains (Fig. 4D). Hippocampal neurons tend to gain H3K79me2 and H3K18/K23ac, which is accompanied by a loss of H3K27me2. In addition, all neuronal populations exhibit reduced levels of modified H3K36 (Fig. 4D). These alterations of the histone modification landscape differ vastly both in trend and amplitude from KO animals.

Finally, the analysis of the linker histone H1 revealed no alterations in total levels across the investigated cell types (Fig. S3F). Variant analysis, however, shows neuron-specific changes in variant abundance and relative contributions (Fig. 4E). This revealed opposing regulation of individual variants in granule layers, resulting in the observed unchanged levels of total H1 in those populations (Fig. 4E, Fig. S3F). Particularly, the DG and Cb KI brains are associated with a loss of replication-independent variant H1.0 and gain of H1.4, while the H1 composition of pyramidal neurons seems largely unaffected by the mutation (Fig. 4E).

## Discussion

RTT-associated mutations of *Mecp2* alter transcriptional regulation by disrupting its ability to both, bind to methylated DNA and interact with co-repressors^4,9,51,67^. Mecp2 has been shown to modulate chromatin structure and function on multiple scales, disruption of which is associated with Rett syndrome^1,10,68^. However, due to the high heterogeneity of utilized model systems, RTT mutations and applied methodologies, the observations are inconsistent and understanding remains incomplete. Particularly the effect of loss-of-function of Mecp2 on histone proteins, the building blocks of chromatin, is largely unexplored. Multiple studies have shown that the composition of histone proteoforms within chromatin plays an important role in determining the epigenetic potential of a cell. Different cell types can be identified by their histone proteoform composition^69^ and changes in this composition is associated with a change in phenotype. Despite clear indications of direct interaction between Mecp2 and nucleosomes as well as the ability of Mecp2 to modulate histone modifications, a comprehensive analysis of the histone proteoform landscape is lacking.

To fill this gap, we used a dual spatial proteomic approach to determine the composition of histone proteoforms in different neuronal populations of WT mice and RTT models. Our approach overcomes key limitations of antibody-based assays, including cross reactivity and reagent availability, by allowing precise quantitative profiling of histone proteoforms with unprecedent multiplexing capacity. The pipeline enabled highly selective profiling of histones by utilizing a relatively low number of pure neuronal cells (30000 – 200000 cells) precisely excised from both healthy and Rett syndrome mice. The results revealed surprising heterogeneity in histone composition across neuronal subtypes, varying both by cell identity - granule cell or pyramidal neuron - and the region of origin - hippocampus or cerebellum (Fig. 5C). Notably, this implicates not only differential deposition of histone variants and modifications but also total core histone levels (Fig. 5C). Changes in total protein levels are not accompanied by proportional changes in modified peptides, suggesting substantial contribution of the unmodified forms (Fig. S2E, S3C). This pattern strongly points to the possible involvement of the non–chromatin-bound histone pool, suggesting dysregulation in histone protein synthesis, turnover, and/or degradation^70,71^. Such regulation, if active in neurons, represents an underappreciated layer of epigenetic control, potentially influencing transcriptional responsiveness and cellular identity. Moreover, increased levels of core histones like H3 and H4 have been shown to reduce nucleosomal accessibility and hence decelerate RNA polymerase II based transcription elongation rates, mechanisms that have been linked to extended cellular lifespan^72^. If altering histone levels trigger similar dynamics in neurons, they may serve not only to modulate transcription but also to influence neuronal longevity, adding another dimension to how histone abundance could shape long-term neural function in RTT.

**Figure 5:**
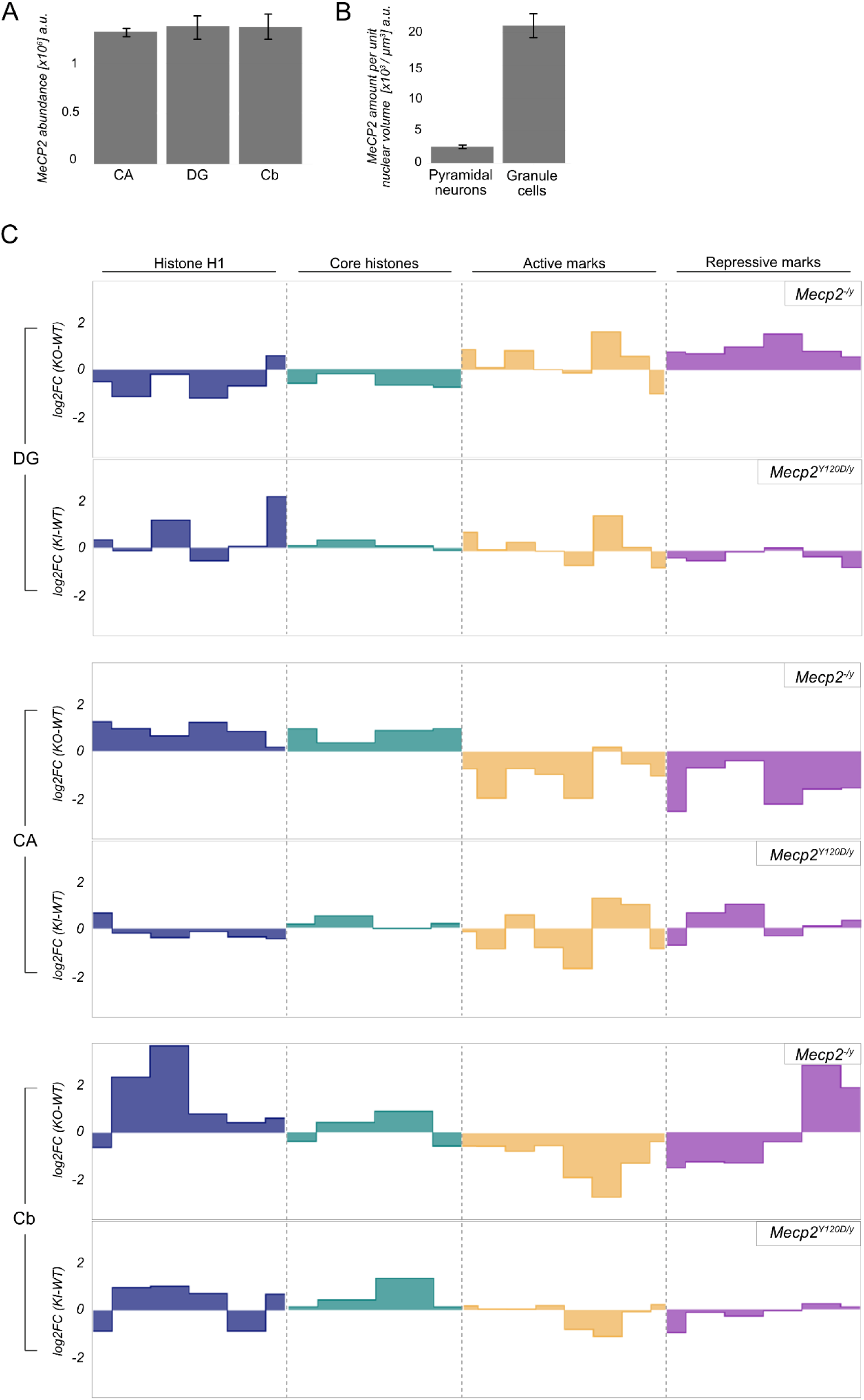
Global shifts in histone landscapes across Mecp2-deficient neurons. **A.** Average abundance of *Mecp2* in WT CA, DG and Cb per nucleus. **B.** Density of *Mecp2* per nuclear volume [µm^3^]. **C.** Area plots illustrating fold changes in linker histone H1 variants (blue), core histones (green) and active (orange)/repressive (pink) post-translational histone modifications in DG, CA, and Cb neuronal populations of *Mecp2^-/y^* (top panels) and *Mecp2^Y120D/y^* (bottom panels) mice compared to WT. Histone proteoforms are ranked along the abscissa by name, each bar represents one proteoform.

The ambiguous effects we observe globally can be explained by the varying local function of Mecp2. It not only maintains a silenced state both by HDAC recruitment and by nucleosome binding and array compaction^6,10^ but also safeguards transcriptionally active nucleosome-free regions from nucleosome invasion^17^. Thus, loss of Mecp2 function can drive divergent chromatin remodeling outcomes, including both destabilization of compacted chromatin as well as compaction of nucleosome-free regions. These structural disruptions can be accompanied by cell type–specific gains or losses of key histone modifications, reflecting bidirectional shifts in chromatin state. Such context dependent alterations likely contribute to widespread transcriptional instability observed in RTT^16,73–75^, where disruption of chromatin states may underlie impaired neuronal function and identity.

Interestingly, despite its high abundance, Mecp2 binding to chromatin has been shown to depend on the nuclear size and state of chromatin compaction^76^. As nuclear size differs vastly between pyramidal neurons (∼10-20 µm diameter) and granule cells (∼5-10 µm diameter)^77,78^, we speculated nuclear Mecp2 concentration to be a determinant in defining the aforementioned neuronal identity. Despite similar total nuclear Mecp2 levels across neuronal types (Fig. 5A), its concentration is significantly higher in the smaller granule cells of the DG and Cb than in CA pyramidal neurons (Fig. 5B), suggesting a molecular basis for the selective neuronal responses. Consequently, the previously observed chromatin alterations within the two mouse models investigated is most closely mimicked by the CA pyramidal neurons^50^. Characterization of the chromatin features in these models was based on cortical neurons, a mixed population that largely consists of pyramidal neurons, suggesting cellular identity to be at least partly influencing the outcomes in RTT. Yet how the complex role of Mecp2 are realized in these different nuclear contexts, is unclear.

Neuronal selectivity is preserved in *Mecp2^Y120D/y^* mice as well. However, the pattern of changes differ markedly from that observed in *Mecp2^−/y^* mice (Fig. 5C, bottom panels). Notably, neuronal populations of *Mecp2^Y120D/y^* exhibit more subtle alterations of histone proteoforms with an overall effect of smaller magnitude (Fig. 5C).

In knock-in mice, the observed alterations primarily stem from disrupted binding of the MBD domain to 5mC/5hmC and impaired recruitment of interactors^50^. In contrast, the Mecp2 *null* mice also include effects caused by loss of non-MBD domains on chromatin, like the AT hook or C-terminal domain, both of which have been implicated in chromatin binding^79–83^. Moreover, complete Mecp2 loss can trigger compensatory responses. These combined effects likely account for the more pronounced chromatin alterations and phenotypic severity observed in the *Mecp2^-/y^* model.

While the link between Mecp2 and core histones is still emerging, its association with linker histone H1 is better established, though not fully understood. Increased H1 levels have been reported as a compensatory response for Mecp2 loss in neurons^50,51^, yet H1.0 does not compete with Mecp2 for binding sites and shows no change in occupancy^21^. By quantifying H1 variants individually, we observe that several – but not all – are elevated in the CA of Mecp2 *null* brains as well as cerebellar granule cells of both *Mecp2^-/y^* and *Mecp2^Y120D/y^*. Notably, the H1 composition differs between the RTT mutants (Fig. 3B, Fig. 4E). The variant specific changes in H1 levels may reconcile previously conflicting findings in the Mecp2 deficient context. These findings, therefore, emphasize the need for detailed investigation into Mecp2’s interaction with replication-dependent H1 variants and their response in its absence.

Our findings underscore the need for spatially resolved analyses of RTT brains, given the intricate, cell-type specific roles of Mecp2. While this study provides novel insights into the global landscape of intact histones and their modified proteoforms, it does not capture localized chromatin dynamics. Future efforts should aim to map these local effects across distinct chromatin compartments and neuronal populations, and to chart their progression over the disease course. Crucially, *Mecp2^-/y^* and *Mecp2^Y120D/y^* models display distinct epigenetic landscapes, reflecting mutation-specific chromatin responses. These differences reinforce the potential importance of applying personalized disease management strategies tailored to the specific Mecp2 mutation involved.

## Data Availability

The mass spectrometry proteomics data have been deposited to the ProteomeXchange Consortium via the PRIDE^84^ partner repository with the dataset identifier PXD065165. Following are the detailed reviewer credentials: Please login using the following token

**Project accession:** PXD065165, **Token:** o3vTwc3BWqF8 or alternatively using the following account details

**Username:**, **Password:** YvX69DqxzKdz

## Acknowledgements

The authors thank Prof. Dr. Changjun Yin, Prof. Dr. Andreas Schober, Claudia Geißler and Dr. Maliheh Nazari Jahantigh from the Institute of Cardiovascular Prevention (IPEK), LMU, for their immense help by providing access to and training for the LMD instrumentation. This work was supported by grants from the Deutsche Forschungsgemeinschaft (DFG) to AI (CRC1123-TPZ02, CRC1309-TPB03, CRC1064-TPZ03). Part of the work was supported by the laboratory funding of Forschungsmodul Medizin program of the Medical Faculty, LMU to SL. We also acknowledge the support of the Italian parents’ association proRETT and of the SCALE-UP project of the Department of Medical Biotechnology and Translational Medicine to NL. The authors would like to thank Dr. Victor Solis of Genedata for his initial assistance with analyzing histone PTMs.

